# Mediator-Microorganism Interaction in Microbial Solar Cell: a Fluo-Electrochemical Insight

**DOI:** 10.1101/2020.03.01.970954

**Authors:** Léna Beauzamy, Jérôme Delacotte, Benjamin Bailleul, Kenya Tanaka, Shuji Nakanishi, Francis-André Wollman, Frédéric Lemaître

## Abstract

Microbial solar cells that mainly rely on the use of photosynthesic organisms are a promising alternative to photovoltaics for solar electricity production. In that way, we propose a new approach involving electrochemistry and fluorescence techniques. The coupled set-up Electro-Pulse-Amplitude-Modulation (“e-PAM”) enables the simultaneous recording of the produced photocurrent and fluorescence signals from the photosynthetic chain. This methodology was validated with a suspension of green alga *Chlamydomonas reinhardtii* in interaction with an exogenous redox mediatior (2,6-dichlorobenzoquinone; DCBQ). The balance between photosynthetic chain events (PSII photochemical yield, quenching) and the extracted electricity can be monitored overtime. More particularly, the non photochemical quenching induced by DCBQ mirrors the photocurrent. This set-up thus helps to distinguish the electron harvesting from some side effects due to quinones in real time. It therefore paves the way for future analyses devoted to the choice of the experimental conditions (redox mediator, photosynthetic organisms…) to find the best electron extraction.

## INTRODUCTION

Over the past ten years, many biophotoelectrochemical systems have been implemented to produce electricity from photosynthesis. They take benefits from a light converter into electricity by notably involving isolated photosystems,^1-4^ thylakoid membranes,^5-8^ or single chloroplasts.^9^ However, in the quest for new energy sources, the use of photosynthetic organisms for their natural ability to capture and use solar light is becoming more and more attractive.^10-13^ High expectations especially concern microbial solar cells, where living photosynthetic microorganisms act as energy converters between light and electricity but are further able to be cultured and self-repaired. The photosynthetic chains are basically composed of a series of redox-active molecules that exchange electrons (see Fig.1A). An alternative electron pathway can be generated to partially “re-route” the photosynthetic electron flux towards an electrode. Accordingly, a key point is the use of an exogenous redox mediator that acts as an electron shuttle and travels back and forth from inside the photosynthetic organisms to the collecting electrode located in the surrounding aqueous solution.^14-18^ However, the electrons originally coming from the photosynthetic chains are embedded in biological membranes and not easily accessible. It therefore raises the question of the best photosynthetic organism/redox mediator tandem.

**Figure 1.**
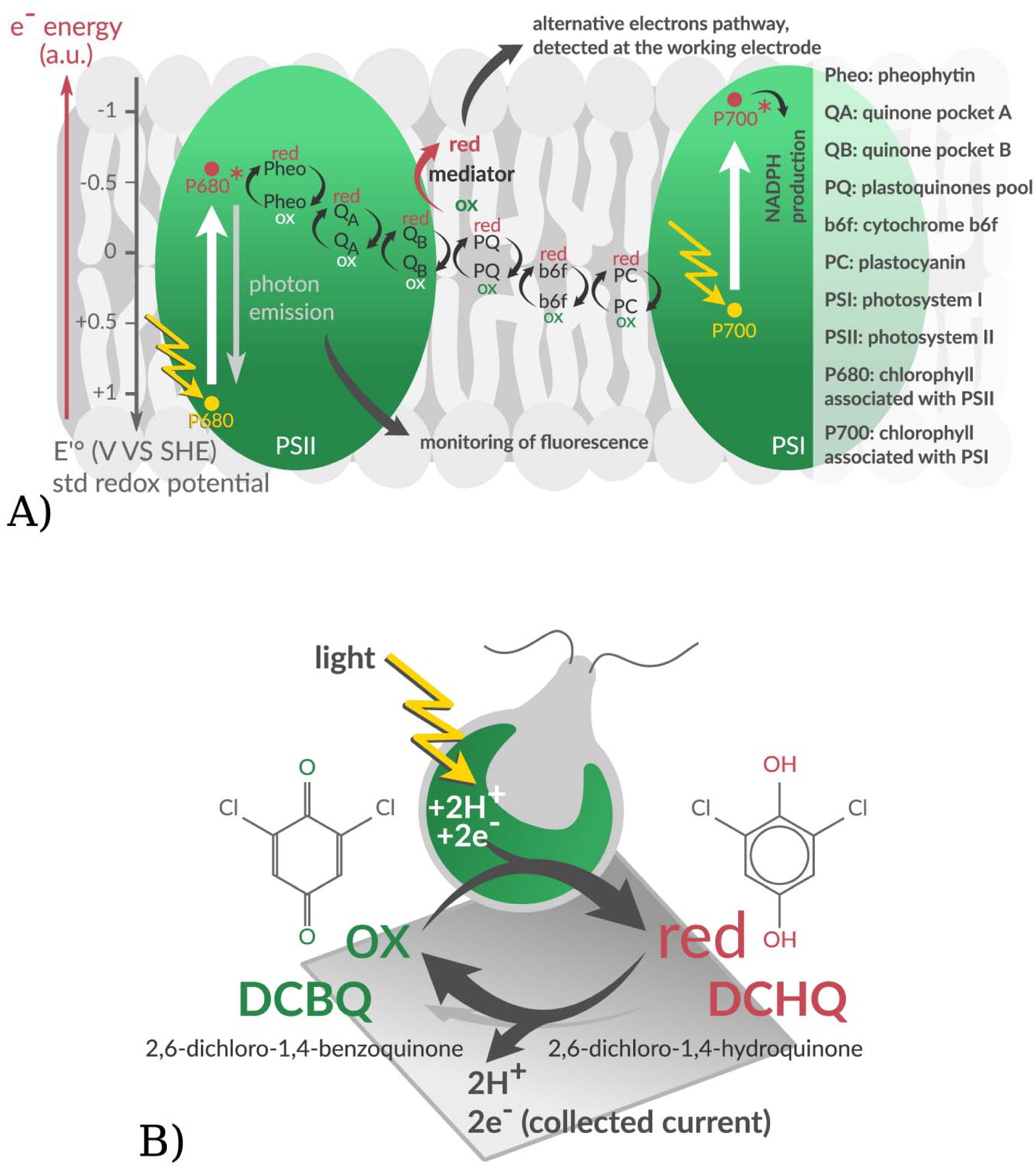
Principle of photosynthetic electrons re-routing by the redox mediator. A) Scheme of the first steps of photosynthesis where the soluble redox mediator can interact with endogenous redox-active molecules embedded into thylakoid membranes (see text). Fluorescence (red/gray arrow pointing downwards) is a deexcitation pathway that can be detected through the Pulse-Amplitude-Modulation (PAM) analysis. B) Electron harvesting from a microalga with an exogenous quinone. The oxidized form (DCBQ) can be reduced (DCHQ) when interacting with the chloroplast of an illuminated alga. DCHQ is oxidized by the working electrode, leading to a measurable current and DCBQ available for a new cycle.

A typical example of model system is the micro-alga *Chlamydomonas reinhardtii*, where the photosynthetic chains are embedded in thylakoid membranes, internal structures of the unique chloroplast. In that case, exogenous quinones can be used, for instance the redox couple 2,6-dichlorobenzoquinone/2,6-dichlorohydroquinone (DCBQ/DCHQ; see Fig.1B).^19,20^ The harvested electrons originally come from the photooxidation of chlorophylls embedded into structured groups of proteins called photosystems I (PSI) or II (PSII). Indeed, exogenous quinones can interact with the photosynthetic chain, especially the quinone Q_A_ after insertion in the pocket B (Q_B_) of PSII and the plastoquinones pool (PQ).^21-24^ The electron harvesting is historically studied by chronoamperometry, an electrochemical technique where the collected current is monitored over time at a polarized electrode. The rise of the collected current is an evidence that the reduced form of the mediator (DCHQ) is produced.^25^ The technique gives a direct estimation of the produced electricity (and indirectly the effect of the redox mediator) but does not provide direct information about the photosynthetic organism. This makes the improvement of microbial solar cells difficult, due to the lack of deep understanding of the overall redox mediator-microorganism interplay. However, fluorometry techniques (monitoring the fluorescence of PSII-associated chlorophylls) have been developed over the past 80 years to study photosynthesis in-vivo. Important parameters can be quantified, such as the photosynthetic yield of the microorganism, an indicator of its physiological state. Since in redox mediator-based solar cells the current eventually drops,^16,18,26^ the improvement of microbial solar cells clearly requires a technique able to provide information on the microorganism ability to perform photosynthesis. The electrical recording therefore should benefit from its coupling with fluorescence measurements. Such combinations are rather scarce and concern photosynthetic biofilms. Together with electrical performances, fluorescence can be used for imaging (confocal microscopy) or to globally indicate photosynthetic activity without further treatment.^27^ An approach combining electrochemistry and Pulse Amplitude Modulation fluorescence (PAM) has been focused on correlations between cell voltage and photosynthetic electron transfer rate in photosynthetic biofilms without redox mediator.^28^ In this context, we report here on a new approach of the electrochemistry/PAM fluorescence combination. The capability of our approach is extended further. First of all, an air-bubbled algal suspension able to maintain the same metabolic state for long periods of time (40 min without any risk of anaerobiosis) was considered. Secondly, a dynamic correlation between electrochemical (photocurrent) and treated fluorescence data (photosynthetic yield; non photochemical quenching) was achieved in real time.

## EXPERIMENTAL SECTION (see details in SI)

### Algae

*Chlamydomonas reinhardtii* line (WT T222+ ecotype) was grown in Tris-Acetate-Phosphate (TAP) medium and resuspended in the exponential phase of growth at 2×10^7^ cells/ml in “Minimum” medium (no carbon source).

### Redox mediator

A 10 mM stock solution of 2,6-dichloro-1,4-benzoquinone (“DCBQ”) was prepared in pure ethanol from the powder version (Sigma-Aldrich), and kept in dark at 4 °C between experiments. During experiments 2/5/10/20 µl of the mother solution were injected into the 2 ml algal suspension for a final concentration of 10/25/50/100 µM, respectively.

### Experimental set-up: Electro-Pulse-Amplitude-Modulation (e-PAM)

An electrochemical cell is designed to be adapted fluorescence measurements with a Pulse-Amplitude-Modulation (PAM) machine (see Fig.2). The working electrode is a transparent square of ITO-coated glass. The reference (Ag/AgCl/KCl sat.) and counter (Pt wire) electrodes dip into the algal suspension. The lights used for excitation and fluorescence measurements are guided by the unique fiber of the PAM-machine below the electrochemical cell. The inner diameter of the glass tube is exactly the same as the one of the fiber (1.1 cm) and defines the area of the working electrode (0.95 cm^2^). The spectroelectrochemical cell is completed with plastic bottom and top parts (yellow in Fig.2A). An air bubbling is also implemented for preventing aerobiosis and sedimentation of the algal suspension.

**Figure 2.**
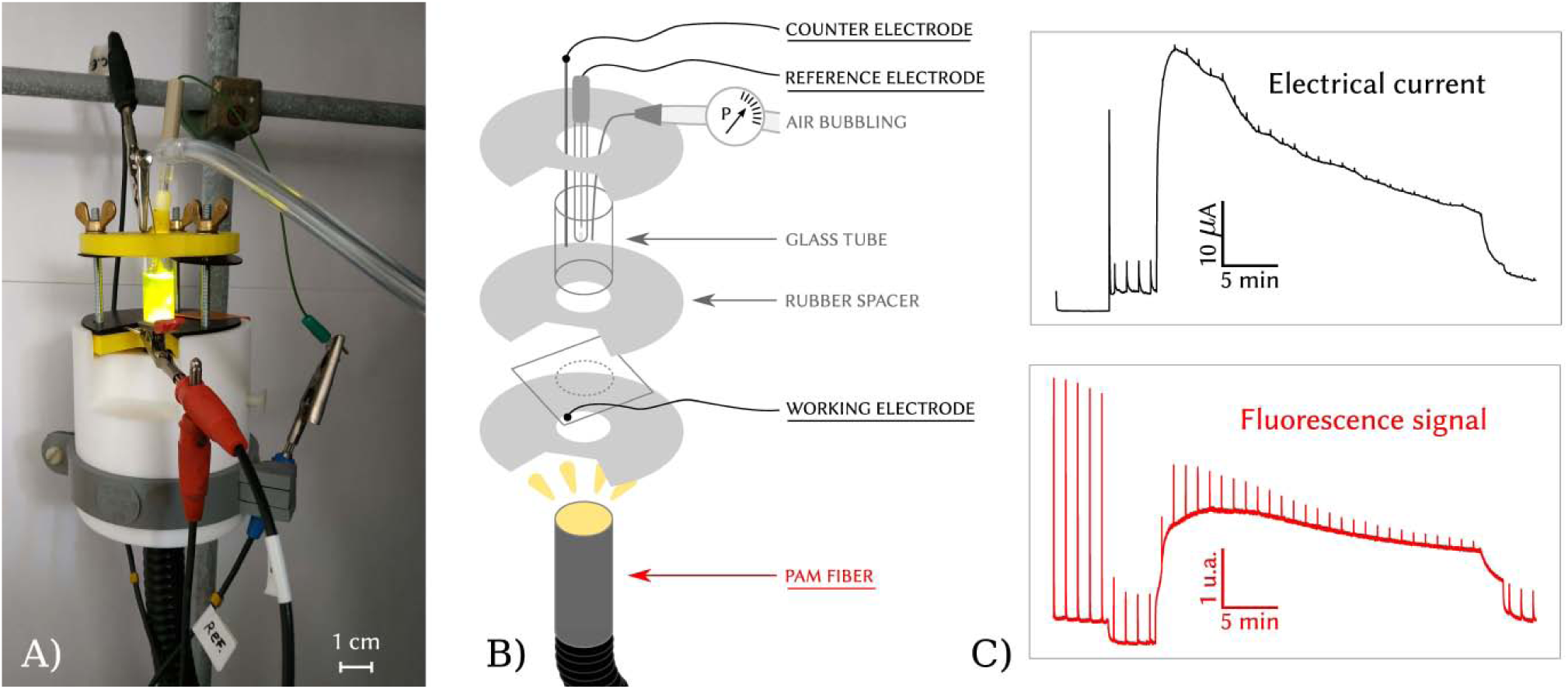
“e-PAM” set-up. A) Picture of the fluoroelectrochemical cell fixed on top of the PAM fiber. B) Corresponding scheme: a glass tube is pressed on top of a ITO-covered glass (working electrode) and define the content of the electrochemical cell in which the algal suspension can be poured. The PAM fiber touches the non-coated side of the glass plate (bottom side). Counter and reference electrodes deep into the algal suspension as well as the air-bubbling needle. C) Examples of raw experiments. Top, black: Electrical current obtained thanks to the electrochemical cell. Bottom, red: Fluorescence signal obtained thanks to the PAM machine.

Chronoamperometric measurements were performed at 0.9V vs Ag/AgCl with the spectroelectrochemical cell at 25 °C by using an Autolab PGSTAT100N potentiostat (Metrohm). The output was digitized at 2 Hz and displayed in real time with Nova 2.0 software with no subsequent digital filtering. Synchronization with fluorescence measurements was achieved by means of an e-corder 821 converter (eDAQ).

Fluorescence measurements were done using a chlorophyll fluorometer PAM101 from Walz, connected to the computer via an e-corder 821 converter (eDAQ). A 3-lights system is used for measuring, actinic and saturating lights and is guided through the unique end of the PAM fiber. A Schott lamp (KL 1500 LCD) is responsible for the white actinic light (700 µmol m^-2^ s^-1^). A red diode (Thorlabs; M625L3; λ = 650 nm; 2700 µmol m^-2^ s^-1^) provides the saturating pulses controlled by the software Chart (stimulator in pulse mode; every minute and being 350 ms-long).

## RESULTS

### Validation of the electrochemical set-up - Redox mediator concentration drives photocurrent intensity

**“**e-PAM” is an electrochemical cell allowing simultaneous fluorescence measurements (Fig.2). Fig.3A displays the 40 min-chronoamperograms obtained for the algal suspension with 4 different concentrations of DCBQ (without their coupled fluorometry data for more clarity). The applied oxidizing potential (t = 0) promotes a capacitive current that drops quickly. DCBQ is then added to these “dark-adapted” samples (4.5 min after the start of the experiment) at 10, 25, 50 µM or 100 µM. This leads to a “dark current” (∼ 6 µA) independent of the DCBQ concentration. This “dark current” was already detected in previous works and may result from interaction with mitochondria or endogenous stored carbohydrates.^13,25,29^ In contrast, the light-induced current (at 8.5 min) depends on the DCBQ concentration. It results from a photoelectrocatalytical cycle involving the illuminated algae/quinone (DCBQ) tandem and the hydroquinone (DCHQ) oxidation at the electrode surface (Fig. 1B; DCBQ reduction into DCHQ by the photosynthetic chain + electrochemical oxidation: DCHQ = DCBQ + 2e^-^ + 2H^+^).^25,26,30^ This corresponds to a re-routing of the photosynthetic electrons at the electrode surface. The maximum photocurrent value increases with DCBQ concentration and is plotted in Fig.3B. For the assayed concentrations, a linear relationship is observed (slope = 0.17 µA µM^-1^). This behavior is consistent with previous works dealing with the same models and ascertains that the electron harvesting is the rate-determining pathway under these conditions.^25,31^ Finally, at the end of experiments, an inhibitor of photosynthesis, DCMU (3- (3,4-Dichlorophenyl)-1,1-dimethylurea) was added,^32-34^ thus leading to a consecutive drop of the collected current (a similar drop is observed if the light is turned off instead; see later). DCMU interrupts the photosynthetic electron transport chain between Q_A_ and Q_B_, demonstrating that the electron harvesting site of the electron shuttle is mainly located at Q_B_ or downstream, as expected for exogenous quinones.^19,35,36^ Control experiments were also performed in absence of DCBQ (see SI) where no light or only saturating pulses/actinic light are used. In both cases, no current was recorded, thus meaning that the electrical current observed with our set-up really comes from the electron harvesting by DCBQ. All in all, these results validate the set-up from an electrochemical point of view.

**Figure 3.**
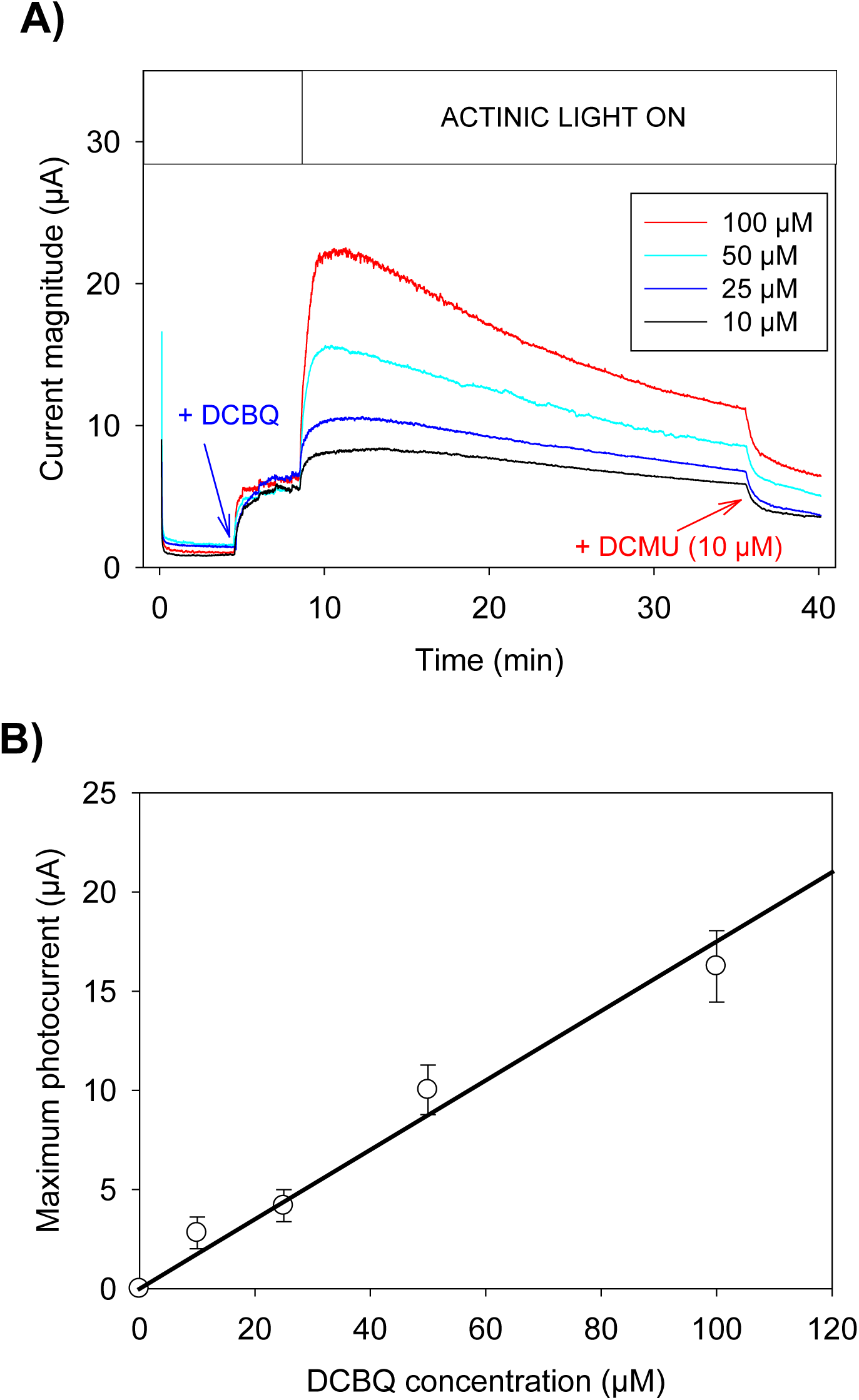
A) 40 min-chronoamperograms obtained for the algal suspension (2×10^7^ cells/ml) with 4 different DCBQ concentrations: 10, 25, 50 and 100 µM. B) Plots of the photocurrent (I_max_ - I_dark_) as a function of DCBQ concentration and its linear fit (y = 0.17×, R^2^ = 0.98).

### First qualitative observations with e-PAM coupling - DCBQ quenches chlorophyll fluorescence during the photocurrent production

Fig.4 displays typical coupled electrochemistry-fluorescence data. Fluorescence measurements rely on Photosystem II (PSII) excitation within the algae suspension. Briefly, actinic light is captured by antenna (LHCII: Light Harvesting Complex II) containing chlorophyll (Chl) whose energy is transferred to the PSII primary donor, P680 (Chl* + P680 → Chl + P680*). It formally leads to a charge separation which results in water oxidation and reduction of the primary acceptor Q_A_ (Fig. 1A) then followed by subsequent electron transfer steps along the photosynthetic chain until the final CO_2_ reduction. Relaxation by fluorescence from excited PSII therefore competes with these electron transfers and depends of the redox state of PSII, i.e. the fraction of open (Q_A_) and closed (Q_A_ ^-^) centers. Of note, another photosystem (PSI) is involved in the photosynthetic chain but contributes little to fluorescence.^37-39^

**Figure 4.**
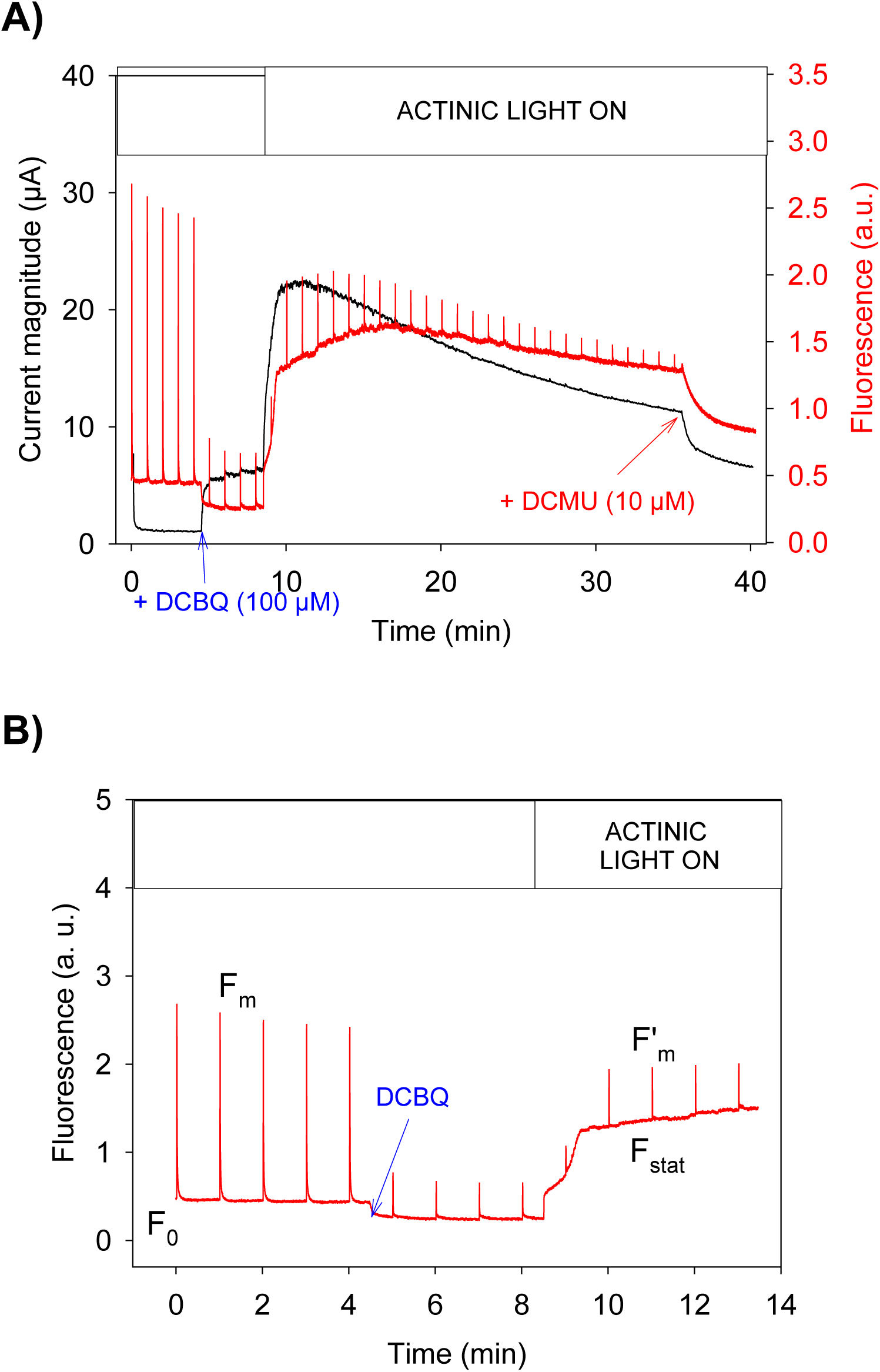
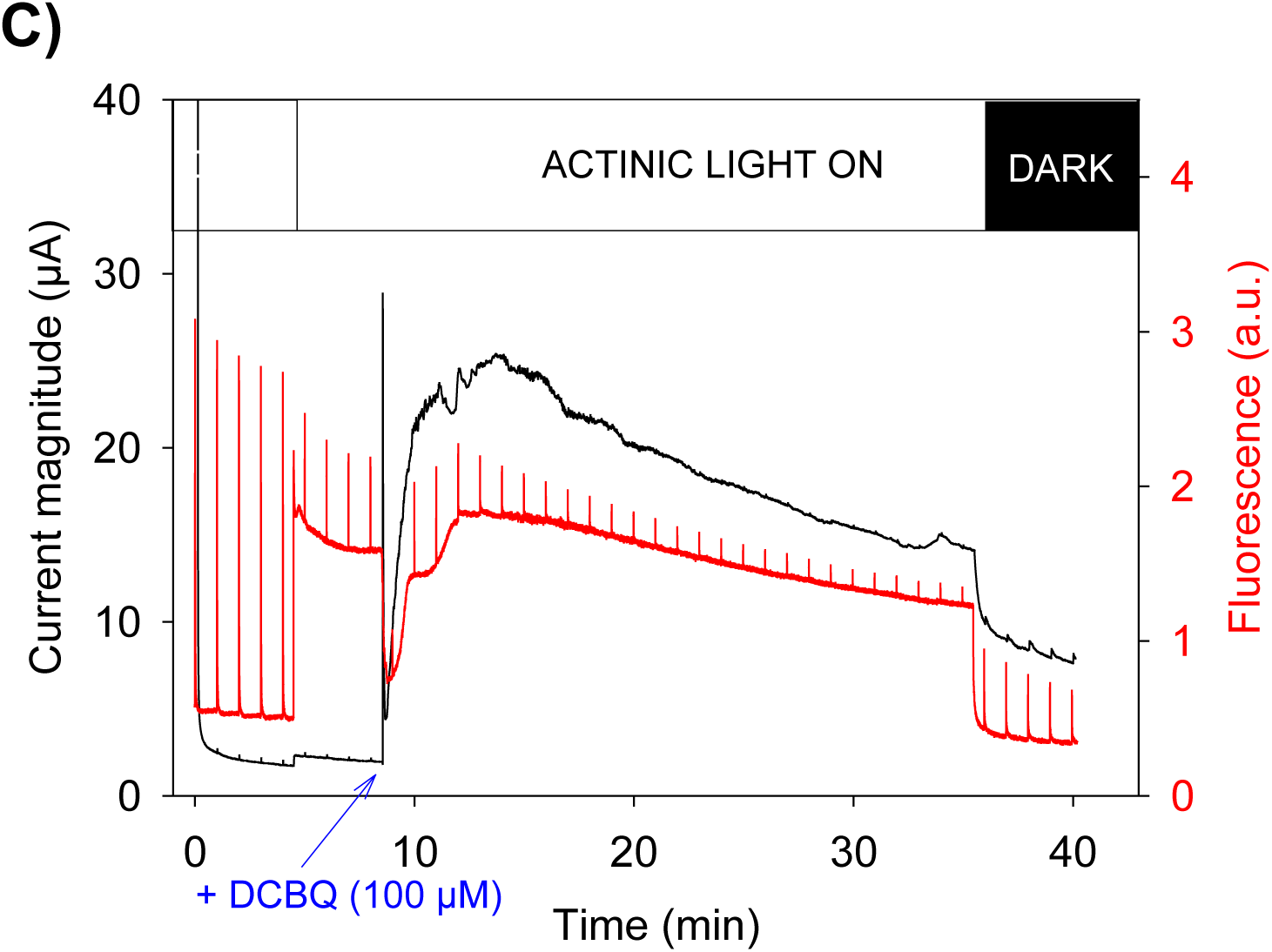
A) Typical data obtained from the coupling of chronoamperometry (black curve, left axis) and fluorescence (red signal, right axis) measurements over time. DCBQ is introduced (blue arrow) before irradiation of the algal suspension. Pulses (350 ms duration) of saturating red light (2700 µmol m^-2^ s^-1^) irradiate the solution every minute. In addition, the actinic white light (700 µmol m^-2^ s^-1^) can be turned on (white bar). B) Zoom of the fluorescence curve that depicts the important fluorescence levels recorded from the algal suspension: in dark without quinone (F_0_); under actinic light (F_stat_) and under the saturating pulse (F_m_ in dark before DCBQ addition and F’_m_ thereafter). C) Typical coupled experiment where DCBQ is added under actinic light. The photosynthetic activity is interrupted by A) adding DCMU or C) turning off the actinic light.

Practically, fluorescence is extracted from different light-conditions (Fig. 4B and SI): under darkness (F_0_), under actinic light (F_stat_) and under a saturating pulse (noted F_m_ or F’_m_ respectively for pulses made before or after DCBQ addition; see below). F_0_ is the minimum fluorescence level when all PSII centers are opened. F_stat_ corresponds to a fluorescence value where the photosynthetic activity occurs with a given PSII photochemical conversion capacity. The saturating pulse is long enough to fully reduce all electron acceptors downstream of PSII, thus closing all the PSII centers (i.e no photochemical conversion capacity for PSII). It leads to the maximum fluorescence level F’_m_. The fraction of photons converted as a photosynthetic activity is therefore proportional to (F’_m_-F_stat_) and help to estimate the photochemical PSII efficiency or yield (Φ_PSII_) defined as (equation (1); see details in SI):

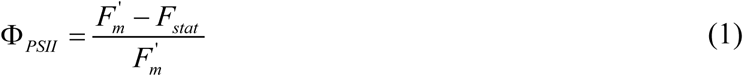

Control experiments with saturating pulses in absence of DCBQ were performed and show that Φ_PSII_ remains quite constant at the timescale of the experiment (see SI).

Two superimpositions of a chronoamperogram and fluorescence measurements are depicted to see the respective effect of the light and the redox mediator (DCBQ is added before (Fig.4A) or after (Fig. 4C) the light is turned on). In the first case a strong decrease of both F_0_ and F’_m_ is observed after DCBQ addition. These changes cannot be explained by electron re-routing. Indeed, in absence of photosynthetic activity in the dark, F_0_ should not be affected. Regarding F’_m_, all reaction centers undergo multiple light-induced charge separations under the light saturating pulse. The photosynthetic chains thus become over-reduced by a flux of electrons which cannot be involved along the photosynthetic chain or counteracted by a DCBQ-mediated re-routing of electrons.^19^ Therefore, this fluorescence level should remain maximal unless an energy dissipation mechanism acts upstream of charge separation in the reaction centers, i.e at the level of light excitation of the antenna pigments. This mechanism is indeed a property that most exogenous quinones exhibit.^19,40-43^ In this case, a direct interaction between the quencher Q and the excited chlorophyll (Chl + light → Chl*) is followed by the formation of a charge transfer complex (Chl* + Q → [Chl*---Q] → [Chl^+^, Q^-^]). In thylakoid membranes, this charge transfer complex then decays to the ground state : [Chl^+^,Q^-^] → Chl + Q).^42,43^

It is worth mentioning that, from a fluorescence point of view, photosynthesis is a process that leads to a quenching of fluorescence. As mentioned in the literature,^37,38^ other pathways (including interaction between exogenous quinones and excited chlorophylls) can contribute to the fluorescence quenching. This is why NPQ (non-photochemical quenching) is defined to reflect the fluorescence decrease related to other pathways than the electron transfer along the photosynthetic chain. Practically, it can be calculated as (equation (2); see details in SI):

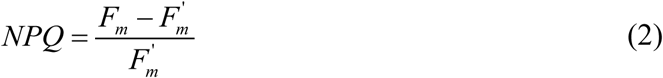

F_m_ is the fluorescence value where no NPQ occurs, i.e. for the last pulse before DCBQ adding in our model system.

When the light is then turned on (Fig. 4A), the NPQ decreases since F’_m_ rises to get closer to the original F_m_ value. Indeed, the oxidized form of the quinone (DCBQ) is a quencher but not its reduced form. Under light, the redox mediator is reduced into DCHQ by the photosynthetic electron flow, leading to the current rise visible on the coupled chronoamperogram. The quenching therefore decreases. In the second experiment (Fig.4C), the actinic light is first turned on and leads to the typical fluorescence rise from F_0_ to F_stat_. The maximum fluorescence value is slightly decreased, showing an endogenous quenching to protect the alga against strong light. When DCBQ is then added, the same phenomenon is observed, i.e. a transient decrease of the F’_m_ value in the first 1-2 minutes, corresponding to the quenching by oxidized DCBQ and quickly after its relaxation as DCBQ becomes reduced by the illuminated algae. After the maximum current, a slow drop is observed, as well as a decrease of both F_stat_ and F’_m_. When photosynthesis is further prevented by addition of DCMU (Fig.4A) or turning off the light (Fig.4C), the remaining current was rapidly reduced to a value close to the dark current. In the presence of DCMU, photosynthetic chains are completely unable to process further any photo-induced electron and F_stat_ should rise to F_’m_. Here F_stat_ merges with F’_m_ as expected, but then decreases, showing again the quenching effect of the oxidized mediator which now accumulates more in the absence of photosynthesis. When the light is simply turned off in Fig.4C, the system goes back to a state similar to what was observed in Fig.4A between 5 and 8 min: the PSII is still active (F_stat_ is significantly lower than F’_m_) and a strong quenching occurs due to almost all redox mediator molecules being back to their oxidized form. 4 and 3 repetitions of each kind were performed with different batch culture of the alga and gave similar results (I_max_ = (33 ± 3) µA). At this stage, these first combined analyses between electrochemistry and fluorescence validate the “e-PAM” coupling.

### NPQ mirrors photocurrent

The non photochemical quenching effect of DCBQ is clearly an important aspect of the alga-quinone interaction that needs to be further analyzed. Indeed, a quinone with a high quenching activity will promote energy losses by indirectly capturing light and will not further be available for the electron re-routing. Using saturating pulses, the overall NPQ (endogenous + exogenous) can be quantified and monitored over time. Fig.5A shows a chronoamperogram and its corresponding NPQ deduced from the fluorescence measurement (from the experiment previously shown in Fig.4A; all replicates show the same behavior). Strikingly, from the moment DCBQ is added slightly before 5 min, the current and the NPQ start to behave in an opposite manner. The experiments done at 4 different DCBQ concentrations (from Fig.3A) helps to estimate the NPQ value just before the light is turned on (t = 8 min). Fig.5B clearly shows that the NPQ is proportional to the DCBQ concentration. This means that the endogenous quenching can be neglected compared to the exogenous one since the overall NPQ can be mostly attributed to the DCBQ alone. The linear relationship also shows that DCBQ is homogeneously distributed in the vicinity of chlorophylls as expected in a homogeneous Stern-Volmer quenching (NPQ = KC_Q_; where K is the quenching constant and C_Q_ the quencher concentration; see details in SI). The quenching constant value for DCBQ is thus equal to (0.027 ± 0.007) L mol^-1^ and is consistent with those found for chloroquinones with *Chlamydomonas reinhardtii* ΔPetA mutants.^19^ The “mirror-effect” between NPQ and photocurrent indicates that the redox changes of the mediator can be tracked in its oxidized form by fluorescence or in its reduced form by electrochemistry. This is a remarkable feature of the “e-PAM” set-up that validates its robustness.

**Figure 5.**
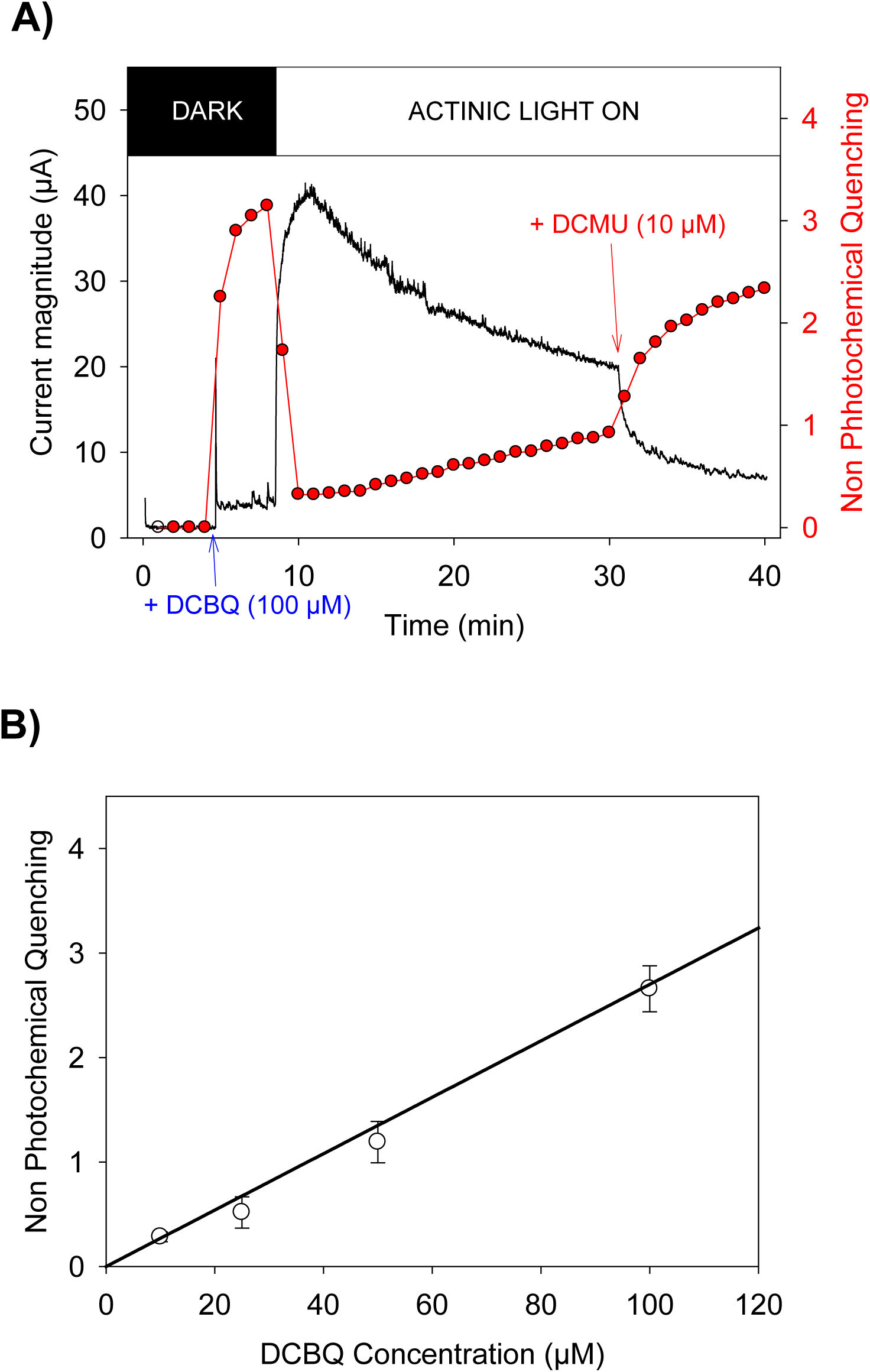
A) Superimposition of a 40 min-chronoamperogram (black curve, left axis) and its corresponding NPQ (red, right axis) calculated every minute from the fluorescence signal. B) Plot of NPQ versus DCBQ concentration (from the experiments shown in Fig 3A; y = 0.027x; R^2^ = 0.98).

### PSII turnover rate is transiently boosted, then collapses

As mentioned above, the photosynthetic yield of PSII (Φ_PSII_) corresponds to the fraction of absorbed light by PSII-associated chlorophylls that ends-up as electrons in photosynthetic chains.^44^ The limiting step of the photosynthetic chain is at the oxidizing site of the b_6_f complex,^45^ forcing PSII to work below its maximum turnover rate. The electron re-routing occuring between PSII and b_6_f is thus expected to increase the PSII turnover rate in presence of an exogenous electron acceptor. The PSII photochemical yield is a relevant data to compare with the photocurrent. Fig. 6 displays the superimposition of a chronoamperogram and Φ_PSII_ (from the experiment in Fig.4B). After its typical drop when the light is turned on (from 0.8 to 0.3), Φ_PSII_ transiently increases after adding DCBQ before decreasing a few minutes later (see zoom in Fig. 6B). Of note, the increase of Φ_PSII_ is especially difficult to observe in our context since the quenching effect of quinones contributes in the opposite way. Despite this limitation, this Φ_PSII_ increase is reproducibly observed (n =8; average increase = (18 ± 4) %). This indicates that the re-routed electrons are - at least partially - “excess electrons”. They result from the use of the excess of absorbed energy that the algae would not process for regular photosynthetic electron transfer, i.e. electrons that rather induce heat dissipation, photon reemission or even photosynthetic damage induced by back-reactions in the reaction centers under saturating light, as it occurs in nature.

**Figure 6.**
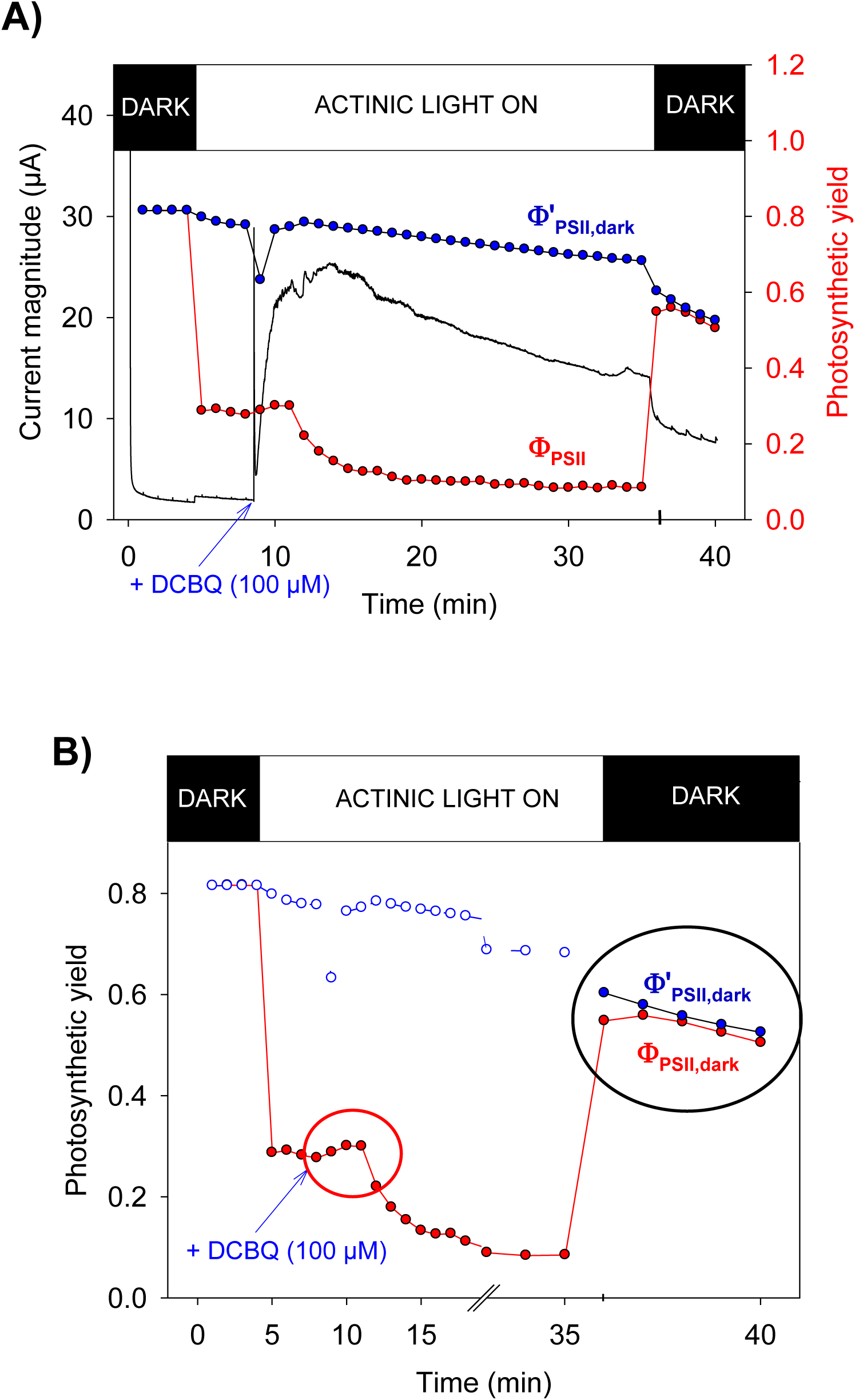
A) Superimposition of a chronoamperogram (black curve, left axis) and its corresponding Φ_PSII_ (red, right axis) calculated every minute from the fluorescence signal. The corrected Φ_PSII_ (Φ’_PSII,dark_; blue) is plotted by taking into account only the quenching effect of DCBQ. This estimated evolution of Φ_PSII_ in the dark under the same level of NPQ has meaning only in the dark (right side of the dotted vertical line). B) Zoom. The Φ_PSII_ increase is observed at short times after applying actinic light. After turning off the light, comparison between the actual Φ_PSII,dark_ and its corrected value Φ’_PSII,dark_ can be made. The irrelevant values in absence of darkness are represented as dashed lines and circles.

Furthermore, in all the experiments, the rapid rise of the current is followed by a slower phase of current decrease (∼ 50% of the current has decreased after 20 min). The reason of this drop remains unclear and the side-effects of quinones are still under debate, but it is known that exogenous quinones tend to be toxic.^46-48^ Therefore the current drop could reflect the progressive accumulation of cellular defects due to quinones or it could also be due to quenching, i.e. a re-routing phenomenon becoming less efficient over time, independently of the physiological state of the algal cells. Interestingly, when Φ_PSII_ was decreasing, the current was still rising on the chronoamperogram, meaning that the DCBQ-induced quenching was not significant at this stage and could therefore not be responsible for the decreased electron flow. This is more likely the signature of DCBQ toxicity, affecting the photosynthetic chain and reducing its kinetics. The evolution of Φ_PSII_ during the 5 first minutes after DCBQ addition thus reflects a complex interplay between rapid changes in electron re-routing, non photochemical quenching and DCBQ toxicity.

### Correlations with “Φ_PSII_ in the dark”

In any event, the electron extraction is more efficient at short times before competitive phenomena take place and lead to the current decrease. To try to disentangle these effects, a quite simple relationship (equation (3)) can be found under dark conditions. It estimated how the maximum Φ_PSII_ (“Φ_PSII_ in the dark; Φ_PSII,dark_) would be theoretically affected by only the quenching effect of DCBQ according to (see SI):

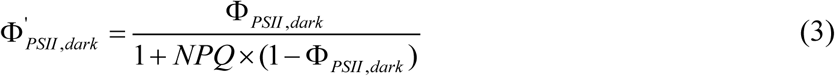

This theoretical Φ_PSII_ (Φ’_PSII,dark_) can be calculated from the corresponding experimental NPQ value and the initial photochemical yield in the dark Φ_PSII,dark_ (taken at the 4th experimental point when there is no actinic light nor DCBQ). The results are plotted in Fig. 6 (zoom in Fig. 6B). Φ’_PSII,dark_ is compared with the experimental Φ_PSII_ for periods of experiment performed in the dark, i.e. when the light was turned off. Strikingly, the two plots globally merge in that time period. Therefore the drop of Φ_PSII_ in the dark observed at the end of the experiment can be attributed mostly to the quenching. On average (n = 5), the quenching can explain (80 ± 4 %) of the drop of Φ_PSII_ in the dark. The DCBQ toxicity which is responsible for the significant drop of Φ_PSII_ (t = 12 min) under light is less visible at the end of the experiment in the dark. This apparent contradiction is explained by the fact that Φ_PSII_ in the dark is a measurement of the maximum PSII turnover rate (not slowed-down by the rest of the photosynthetic chain). Potential defects downstream PSII are not visible in these conditions. Moreover, the measurement of Φ_PSII_ in the light yields the turnover rate of the all photosynthetic chain, i.e. the turnover rate of its limiting step. This therefore suggests that DCBQ-toxicity probably affects a redox intermediate downstream PSII, but not the PSII itself.

## CONCLUSION AND OUTLOOK

We built a fluo-electrochemical set-up (e-PAM) that simultaneously monitors the photosynthetic chain and the redox mediator behavior when the two are exposed to each other. The validation of this analytical combination has been demonstrated through analyses with a model system involving an algal suspension of *Chlamydomonas reinhardtii* and an exogenous quinone (DCBQ) as a redox mediator. The oxidized mediator showed quenching properties that significantly affect the fluorescence of the microorganism, making the molecule traceable in its oxidized form by fluorometry as well as in its reduced form by chronoamperometry. This “e-PAM” set-up described here is thus able to quantify the non photochemical quenching, the PSII photochemical yield and photocurrent. During the first minutes after its addition, the redox mediator transiently boosts the PSII yield. Such a Φ_PSII_ increase stands as a proof of concept that the excess energy that photosynthetic organisms absorb under high light can be extracted by not compromising their vital photosynthetic activity at short times. Furthermore, a correlation between drop of current and rise of non photochemical quenching by the quinone was observed. This further validates the set-up but also opens the question of a potential “vicious circle” effect of the quenching properties of DCBQ. Whatever the original reason for the decrease of the re-reduction rate by algae, the oxidized form prevents the photosynthetic chains to have access to the light. This therefore reduces their ability to reduce the redox mediator. Independently of a possible toxicity, future choices of redox mediator molecules need to consider quenching properties. This “e-PAM” set-up is expected to be extended to other photosynthetic organisms (other algae, cyanobacteria…) and electron shuttles thus making this approach very promising for future clean energy production and to find new redox mediator molecules with less toxic and very little quenching properties.

## ASSOCIATED CONTENT

The supporting information is avalaible free of charge.

Experimental details; principles of PAM fluorescence measurements and mathematical equations (SI.pdf)

3D structures of top and bottom parts of the electrochemical cell for 3D printing (top_and_bottom.stl)

## ACKNOWLEDGEMENT

This work has been supported in part by CNRS (UMR 8640, UMR7141), Ecole Normale Supérieure, French Ministry of Research, Faculté des Sciences et Ingénierie - Sorbonne Université, and the “Initiative d’Excellence” program from the French State (Grant “DYNAMO”, ANR-11-LABX-0011-01). We are very grateful to Anja Krieger-Liszkay for giving us a free access to her PAM machine. We thank Pr. Pierre Joliot for helpful discussions.

