## supplemental for "Mediator-Microorganism Interaction in Microbial Solar Cell: a Fluo-Electrochemical Insight"

#### Table of Contents


1. **Detailed Experimental Section**
    a. **Algae**
    b. **Redox mediator**
    c. **Experimental set-up: Electro-Pulse-Amplitude-Modulation (e-PAM)**
2. **Fluorescence Measurements – Principles and Equations**
    a. **Principles**
    b. **Photochemical yield**
    c. **The 3-lights system**
    d. **Non photochemical quenching**
    e. **Homogeneous Stern-Volmer quenching**
    f. **Estimation of $\Phi_{PSII}$ in the dark for a given NPQ level**
3. **Supplementary data : control experiments and replicates**

### 1. DETAILED EXPERIMENTAL SECTION

#### a. Algae

*Chlamydomonas reinhardtii* line (WT T222+ ecotype) was grown in Tris-Acetate-Phosphate (TAP) medium (Tris base (20 mM), $NH_4Cl$ (7 mM), $MgSO_4$ (0.83 mM), $CaCl_2$ (0.45 mM), $K_2HPO_4$ (1.65 mM), $KH_2PO_4$ (1.05 mM), $CH_3CO_2H$ (0.3 mM) and resuspended in the exponential phase of growth at $2x10^7$ cells/ml in “Minimum” medium (no carbon source). Before an experiment, algae were gently aerated during 2 hours using an orbital shaker, under white light (photon flux: 8 µmol $m^{-2}$ $s^{-1}$, same as for the growth conditions). The light was turned off 20 min prior to the measurement for the algae to be in a “dark-adapted” state. 2 ml of the algal suspension were then transfered into the electrochemical cell. During the measurements the algae were air-bubbled with a micro-needle, a convenient way to keep them aerobic in homogenous resuspension.

#### b. Redox mediator

We used 2,6-dichloro-1,4-benzoquinone (shortly called “DCBQ”) as an electron carrier. DCBQ is the oxidized form of the molecule, which can be reduced by addition of 2 electrons and 2 protons (see Fig.1B). The reduced form is called 2,6-dichloro-1,4-hydroquinone, or DCHQ, and can be re-oxidized at the surface of a polarized electrode, giving rise to a measurable current. The applied potential on the working electrode to easily oxidize DCHQ molecules is equal to 0.9V vs Ag/AgCl. It was deduced from cyclic voltammograms at ITO surface with DCBQ. The redox potential of the DCBQ/DCHQ couple was found equal to 0.4V vs Ag/AgCl (at pH 7.4) but due to slow kinetics at ITO, a value higher than the oxidation peak (but lower than the water oxidation wall) was chosen to ensure an electrochemical process rate-determined by mass transfer. No photocurrent was observed in a control experiment with illuminated algae alone (see below). It therefore shows that the applied potential did not induce side reactions. Practically, a 10 mM stock solution of DCBQ was prepared in pure ethanol from the powder version (Sigma-Aldrich), and kept in dark at 4 °C between experiments. During experiments 2/5/10/20 µl of the mother solution were injected into the 2 ml algal suspension for a final concentration of 10/25/50/100 µM, respectively.

#### c. Experimental set-up: Electro-Pulse-Amplitude-Modulation (e-PAM)

The set-up is comprised of an electrochemical cell designed to be also convenient for fluorescence measurements with a Pulse-Amplitude-Modulation (PAM) machine (see Fig.2). In this respect the working electrode is a transparent square of ITO-coated glass, which allows proper illumination. The reference (Ag/AgCl/KCl sat.) and counter (Pt wire) electrodes dip into the algal suspension from above to avoid perturbation of the light path. The various lights used for excitation as well as the algal fluorescence are guided by the unique fiber of the PAM-machine that touches the electrochemical cell from below. We choose the inner diameter of the glass tube to be exactly the same as the one of the fiber (1.1 cm): its size defines the area of the working electrode (0.95 $cm^2$) that will be in contact with the algae. This allows the algal population studied by electrochemistry to be as close as possible to the one monitored by the PAM. The rigidity of the structure of this spectro-electrochemical cell is ensured thanks to plastic bottom and top parts (visible in yellow in the picture Fig.2A) that we 3D-printed (corresponding .std files available in the SI). An air bubbling is also implemented for preventing aerobiosis and sedimentation of the algal suspension. It is worth mentioning it leads to experiments under forced convection. In that way, all the suspension is involved in the photoelectrocatalysis responsible for the photocurrent and the fluorescence measurements as well.

##### Electrochemistry

Chronoamperometric measurements at constant potential (0.9V vs Ag/AgCl) were performed with the spectroelectrochemical set-up described above. All the measurements were carried out at 25 °C. The control of the applied potential value and the acquisition of the current-time curves were achieved by using an Autolab PGSTAT100N potentiostat (Metrohm). The output was digitized at 2 Hz and displayed in real time with Nova 2.0 software with no subsequent digital filtering. Synchronization with fluorescence measurements was achieved by means of an e-corder 821 converter (eDAQ).

##### Fluorescence measurements (PAM machine)

Fluorescence measurements were done using a chlorophyll fluorometer PAM101 from Walz, connected to the computer via an e-corder 821 converter (eDAQ). We set the PAM machine with the following parameters: frequency = 1.6 kHz; light int. = 12; gain = 8; and damping = 9. Fluorescence measurements allow one to record important photosynthetic parameters, such as the PSII yield ($\Phi_{PSII}$) and non photochemical quenching (NPQ). To do so, a 3-lights system is required: a measuring light, an actinic light and a saturating light. In the present set-up all these lights are guided through the unique end of the PAM fiber that touches the bottom of the electrochemical cell (with a bit of optic grease in between). The measuring light and the lock-in fluorescence detection are provided by the PAM machine. A Schott lamp (KL 1500 LCD) is responsible for the white actinic light, illuminating the bottom of the electrochemical cell with a photon flux we measured at 700 µmol $m^{-2}$ $s^{-1}$. Finally, a red diode from Thorlabs (M625L3) provides the saturating pulses ($\lambda$ = 650 nm). It is controlled by the software Chart (stimulator in pulse mode), for turning on every minute, and being 350 ms-long. The diode exhibits a photon flux of 2700 µmol $m^{-2}$ $s^{-1}$ at the bottom of the electrochemical cell. Throughout the experiments, the algae content of the electrochemical cell is exposed to different light conditions and fluoresces accordingly: $F_0$ in the dark, $F_{stat}$ under actinic light, and $F_m$ or $F'_m$ under a saturating pulse. Details about fluorescence measurements are given within the text and especially in Supporting Information that includes detailed explanations and demonstrations about the equations involved in measurements of photosynthesis

#### 2. FLUORESCENCE MEASUREMENTS – PRINCIPLES AND EQUATIONS

Note: In all the work we use a model called “Lake Model”. Other models exist - depending on how the interaction between chlorophyll antennae and photosynthetic electron transport chains is modeled - but are less popular in the field. The most important parameter, $\Phi_{PSII}$ (see later), is anyways very robust because its expression as a function of the fluorescence intensities does not depend on the chosen model. The technique of photosynthesis study by fluorescence measurements was first introduced in 1931 by Kautsky. Since then it has been extensively used and improved, and is now a well established technique[1–5].

#### a. Principles

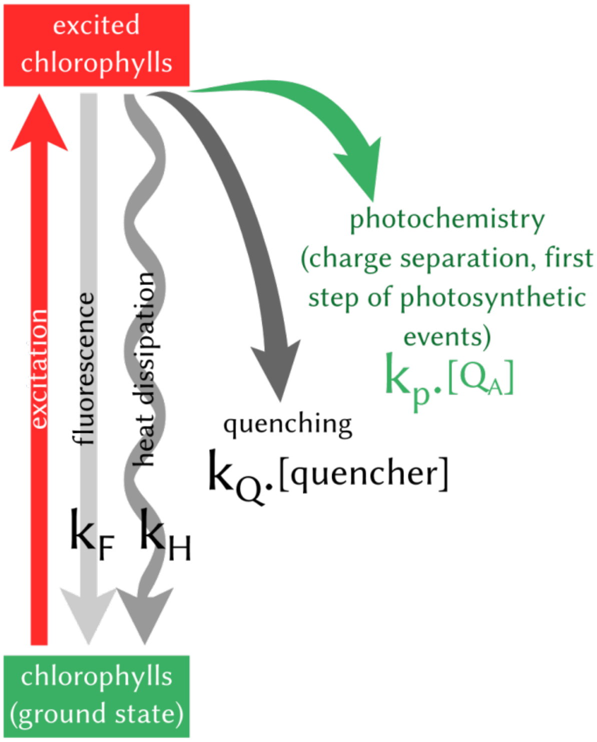


**Figure S1**: Example of deexcitation pathways from the excited PSII-associated chlorophylls. Fluorescence, heat dissipation, quenching and photochemistry are competing and characterized by their rate constants, $k_F$, $k_H$, $k_Q$ and $k_P$ respectively. [$Q_A$] is the proportion of "open centers" (photosynthetic chains with their $Q_A$ molecules in the oxidized form, therefore able to accept an electron). [quencher] is the concentration of the endogenous or exogenous molecules responsible for the quenching. In our case we neglect the endogenous quenching and the only quencher is therefore the exogenous DCBQ.

The principle of fluorescence measurements relies on the simple fact that fluorescence is one of the deexcitation pathways that compete with photosynthetic events. The fate of a photon absorbed by a chlorophyll indeed depends on the rate constants of the different deexcitation pathways: fluorescence (photon re-emission), heat dissipation, quenching (a specific case of heat dissipation that requires the presence of a quencher molecule), or photochemistry (see **Figure S1**). Photochemistry is possible when the delocalized exciton is ultimately transfered to a specific kind of chlorophylls, called P680. The excited P680* transfers an electron to $Q_A$, promoting a charge separation P680$^+Q_A^-$, first step of the photosynthetic electron transfer (a first electron carrier named Pheophytin is in between P680 and $Q_A$, but its short lifetime makes $Q_A$ the "real" first acceptor). If the photosynthetic chain is altered, for example by a chemical, the photochemistry efficiency will drop and the excess energy will be distributed into the remaining deexcitation pathways. The fluorescence will then increase. Its intensity is proportional to the probability of radiative deexcitation, also proportional to the ratio of the fluorescence rate constant to the sum of the reaction rate constants of all competing processes:

$$F \propto \frac{k_F}{\sum k} \quad (1)$$

**b. Photochemical Yield ($\Phi_{PSII}$)**

P680 and $Q_A$ are part of the protein complex PSII, often called reaction center, because responsible for photochemistry. Similarly to the fluorescence efficiency, the photochemical efficiency (or photochemical yield, $\Phi_{PSII}$) is:

$$\Phi_{PSII} = \frac{k_P[Q_A]}{\sum k} \quad (2)$$

where $k_P$ is the rate constant of photosynthesis, and $[Q_A]$ the proportion of PSII with $Q_A$ in its oxidized form. In the dark this proportion is assumed to be 1, and the reaction centers (PSII) are "open", i.e. ready to perform a charge separation. On the contrary, under a saturating light all PSII have $Q_A$ in the reduced form ($Q_A^-$), and $[Q_A]$ is zero. The reaction centers are "closed". Under a typical moderate light that activates photosynthesis without saturating it - such light is called actinic -, $[Q_A]$ ranges from 0 to 1. It is possible to express the $\Phi_{PSII}$ of a sample under an actinic light, as a function of the only measurable values we have access to: the fluorescence intensities. Only 2 measurements are needed: the fluorescence level under the specific actinic light intensity we are interested in (noted $F_{stat}$) and the fluorescence level when a saturating pulse is applied to the same sample (noted $F'_m$) :

$$\Phi_{PSII} = \frac{F_m^{'} - F_{stat}}{F_m^{'}} \quad (3)$$

This relationship results from the fact that the fluorescence intensity is proportional to its efficiency. Under an actinic light we can write:

$$F_{stat} \propto \frac{k_F}{k_F + k_H + k_P[Q_A] + k_Q[DCBQ]} \tag{4}$$

Under a saturating light, $[Q_A] = 0$ and we have:

$$F_m^{'} \propto \frac{k_F}{k_F + k_H + k_Q[DCBQ]} \tag{5}$$

When $k_F$, $k_H$ and $k_Q$ $[DCBQ]$ are constant, we therefore can obtain (using for convenience the notation " $k_{FHQ}$ " *f*or the sum ($k_F + k_H + k_Q$ $[DCBQ]$)) :

$$\frac{F_m^{'} - F_{stat}}{F_m^{'}} = \left( \frac{k_F}{k_{FHQ}} - \frac{k_F}{k_{FHQ} + k_P[Q_A]} \right) \frac{k_{FHQ}}{k_F} \tag{6}$$

$$\frac{F_m^{'} - F_{stat}}{F_m^{'}} = \left( \frac{k_{FHQ} + k_P[Q_A] - k_{FHQ}}{k_{FHQ}(k_{FHQ} + k_P[Q_A])} \right) k_{FHQ} \tag{7}$$

$$\frac{F_m^{'} - F_{stat}}{F_m^{'}} = \frac{k_P[Q_A]}{k_{FHQ}(k_{FHQ} + k_P[Q_A])} \tag{8}$$

$$\frac{F_m^{'} - F_{stat}}{F_m^{'}} = \Phi_{PSII} \quad (according\ to\ eq.\,(3)) \tag{8}$$

In our experiments the DCBQ concentration varies over time, since a variable proportion is reduced into DCHQ after interacting with photosynthetic chains. It would be wrong to determine the initial $F'_m$ value value once and then use it for determining different $\Phi_{PSII}$ during the experiment. This is why we apply a saturating pulse every minute: to keep monitoring the $F'_m$ value throughout the experiment. Therefore, $\Phi_{PSII}$ is calculated each minute by using this instant $F'_m$ and the corresponding $F_{stat}$ that is obtained at the time that just precedes the saturating pulse.

##### c. The 3-lights system

The previous equation (6) is only valid if the factor of proportionality (noted "$\propto$" in equations (1), (4) and (5)) between the measured fluorescence intensity and the efficiency of the fluorescence process - in terms of rate constants - is the same under actinic light and under saturating light. This "∝" factor depends on the set-up instrument and on the excitation light intensity. The last one obviously varies between experiments under actinic or saturating light. This is why a third light source is required, called the measuring light (see **Figure S2**). In Pulse-Amplitude-Modulation (PAM) machines this light source is modulated at a given frequency, and by a system of lock-in amplifier only the corresponding fluorescence (modulated at the same frequency) is collected. This synchronized detected fluorescence is therefore the result of an excitation light that has always the same intensity, so we can compare the fluorescence level over time, when the sample is placed in different light environments. In the dark only the measuring light is on, and its intensity is low enough not to reduce $Q_A$. The photochemistry is therefore favored and the fluorescence is minimal, called $F_0$. Under actinic light the sample is performing photosynthesis: some reaction centers are closed, others are open. Because a fraction of reaction centers is now closed, the photons of the measuring light cannot be converted into photochemistry as efficiently as in the dark, and the remaining fluorescence increases at a level called $F_{stat}$. Under a saturating pulse, all centers are closed. The photons of the measuring light cannot be converted into photochemistry at all and the fluorescence is maximum, called $F'_m$.

In the "e-PAM" coupled set-up the measuring light, as well as the detection of the synchronized fluorescence, are provided by the PAM-machine, guided by an unique "back and forth" fiber (with 2 lateral guides for connecting the other lights, the actinic lamp and the saturating diode) from the bottom of the electrochemical cell to the machine.

Note that more recent PAM-machines are all-in-one and provide the measuring, the actinic as well as the saturating light in one box. The coupled set-up described in this work can be easily adapted to these newest PAM machines.

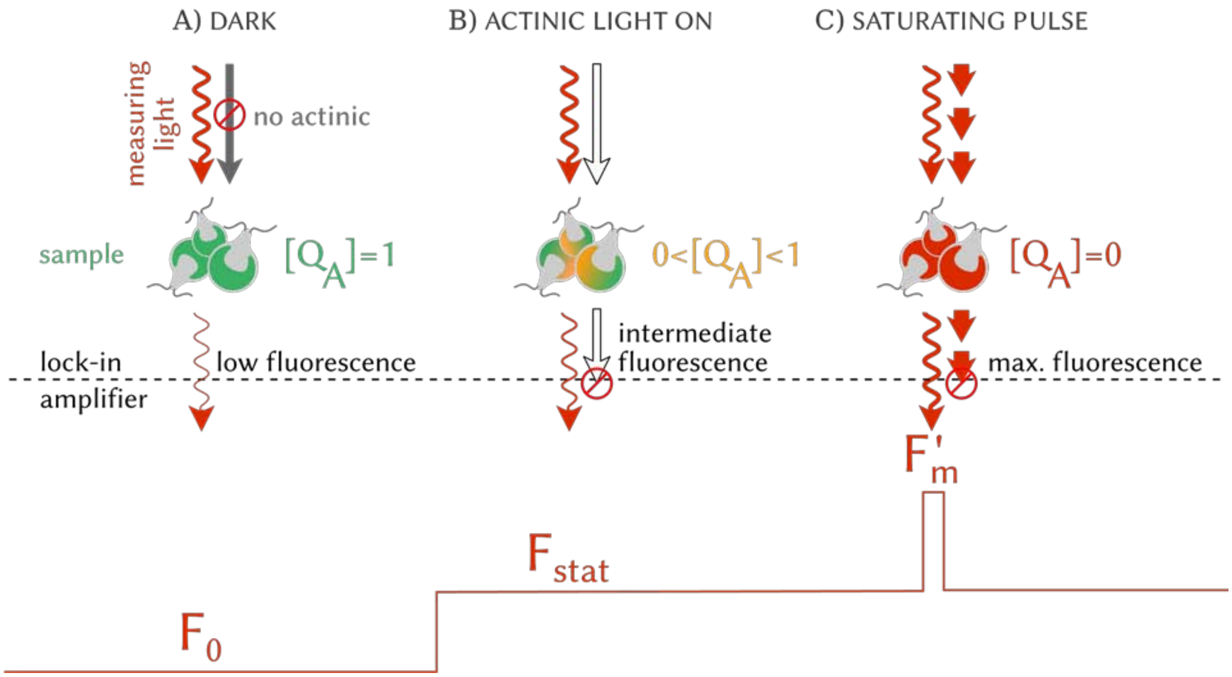


**Figure S2.** Principle of PAM measurements. A) The sample is in darkness, except for the measuring light that is low intensity and modulated. As all $Q_A$ are oxidized, the fluorescence is minimum. The detected modulated fluorescence is called $F_0$. B) The sample is under actinic light (continuous light of moderate intensity), which partially reduces $Q_A$. The fluorescence level is therefore intermediate. The lock-in system filters the fluorescence and collects only the modulated one, called $F_{stat}$. C) The sample undergoes a short but very-high intensity pulse of light (duration = 350 ms, photon flux = 2700 µmol $m^{-2}\,s^{-1}$), called saturating pulse. The intensity of light is chosen to be enough to instantaneously reduce all $Q_A$. No photochemistry is possible, and the detected modulated fluorescence is maximum and called $F'_m$.

##### d. Non-Photochemical Quenching (NPQ)

The non-photochemical quenching or NPQ often refers to the endogenous quenching observed when photosynthetic organisms are exposed to high light. Here the more general definition is considered: any process decreasing the chlorophyll fluorescence when all reaction centers are closed, i.e. reducing $F'_m$ . Typically, an exogenous molecule such as DCBQ, able to accumulate in the vicinity of the chlorophylls and to interact with their excited state to facilitate their relaxation, will promote NPQ. This quencher decreases the photochemical and fluorescence yields by offering a new deexcitation pathway for the excited chlorophylls. From $F_m$ - the maximum reference value in absence of quenching -, the fluorescence under a saturating pulse will drop to $F'_m$ .The mathematical expression of NPQ is therefore expressed as:

$$NPQ = \frac{F_m - F_m^{'}}{F_m^{'}} \qquad (10)$$

$F_m$ is the fluorescence value under a saturating pulse when there is no quenching (in our experiments this value is considered at the last pulse before DCBQ injection) and $F'_m$ corresponds to the fluorescence under a saturating pulse in presence of the quencher (after DCBQ addition).

**e. Homogeneous Stern-Volmer Quenching**

The efficiency of the quenching deexcitation pathway was described as being proportional to the "$k_Q[DCBQ]$" term (see **Figure S1**). Actually this is only valid in the case of a homogeneous Stern-Volmer quenching, i.e. a case where the quencher is homogeneously distributed in the vicinity of the chlorophylls. If so, the previous NPQ expression (eq.10) can be rewritten as a function of the rate constants. $F_m$ is the fluorescence under a saturating pulse before adding DCBQ, so when [DCBQ] = 0, eq.(5) becomes:

$$F_m \propto \frac{k_F}{k_F + k_H} \qquad (11)$$

After DCBQ addition, equation (5) applies :

$$F_m^{'} \propto \frac{k_F}{k_F + k_H + k_Q[DCBQ]} \qquad (5)$$

Therefore the NPQ expression (eq. (10)) becomes:

$$NPQ = \left( \frac{k_F}{k_F + k_H} - \frac{k_F}{k_F + k_H + k_Q[DCBQ]} \right) \frac{k_F + k_H + k_Q[DCBQ]}{k_F} \qquad (12)$$

$$NPQ = \frac{k_Q}{k_F + k_H}[DCBQ] \qquad (13)$$

Under the homogeneous Stern-Volmer quenching assumption the NPQ value is therefore expected to be proportional to the DCBQ concentration.

#### f. Estimation of $\Phi_{PSII}$ in the dark for a given NPQ level

In our experiments the efficiency of photochemistry (i.e. $\Phi_{PSII}$) is affected by the presence of DCBQ. Potentially increased by the re-routing effect of the mediator, the $\Phi_{PSII}$ value is unexpectedly decreased by the toxicity and the quenching properties of DCBQ. To disentangle these effects, we considered the $\Phi_{PSII}$ in the dark, where there is no re-routing. The quenching, acting upstream PSII at the antennae level, affects both $F_0$ and $F'_m$ values. On the contrary, the toxicity *a priori* does not affect the antennae (we neglect this possibility in the present study). The $F'_m$ value (obtained when the photochemistry is saturated) should therefore not be affected by toxicity, even if the photosynthetic chain is damaged, reducing its kinetics. NPQ is therefore a measurement of the quenching alone, unaffected by a potential toxicity. We used the experimental NPQ to estimate how the initial $\Phi_{PSII}$ value (taken in the dark at the beginning of the experiment when there is no light nor DCBQ) would be reduced if the only difference was the presence of quenching.

$\Phi'_{PSII,dark}$ is the expected photochemical yield in the dark in the presence of DCBQ. $\Phi_{PSII,dark}$ corresponds to the initial photochemical yield in the dark. This relationship is only true for a photochemical yield considered in the dark, i.e. when the fraction of open centers $[Q_A]$ is equal to 1. Photochemistry is one of the possible deexcitation pathways, so we can write its efficiency the same way we did previously:

$$\Phi_{PSII,dark} = \frac{k_P}{k_F + k_H + k_P} \quad (14)$$

$$\Phi'_{PSII,dark} = \frac{k_P}{k_F + k_H + k_P + k_Q[DCBQ]} \quad (15)$$

By combining equations (13), (14) and (15), we deduce the relationship used for the data treatment:

$$\Phi'_{PSII,dark} = \frac{\Phi_{PSII,dark}}{1 + NPQ(1 - \Phi_{PSII,dark})} \quad (16)$$

### 3. SUPPLEMENTARY DATA: CONTROL EXPERIMENTS AND REPLICATES

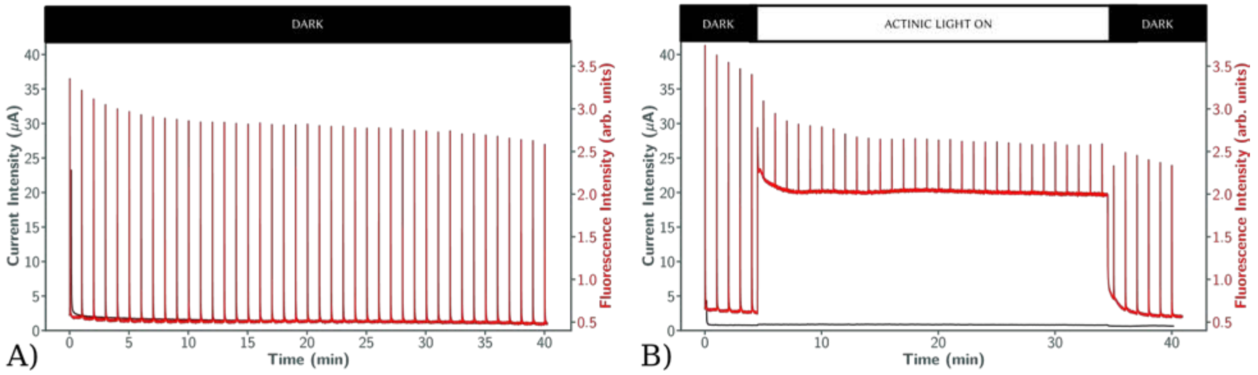


**Figure S3.** "e-PAM" control measurements without DCBQ. A) The algae suspension is in darkness for 40 min, only submitted to the saturating pulses. B) Additionally to the saturating pulses, the algae suspension is submitted to actinic light from 4.5 min to 34.5 min. In both cases, after the rapid decrease of the capacitive current during the first minute, the electrical current then stabilizes at a value close to zero.

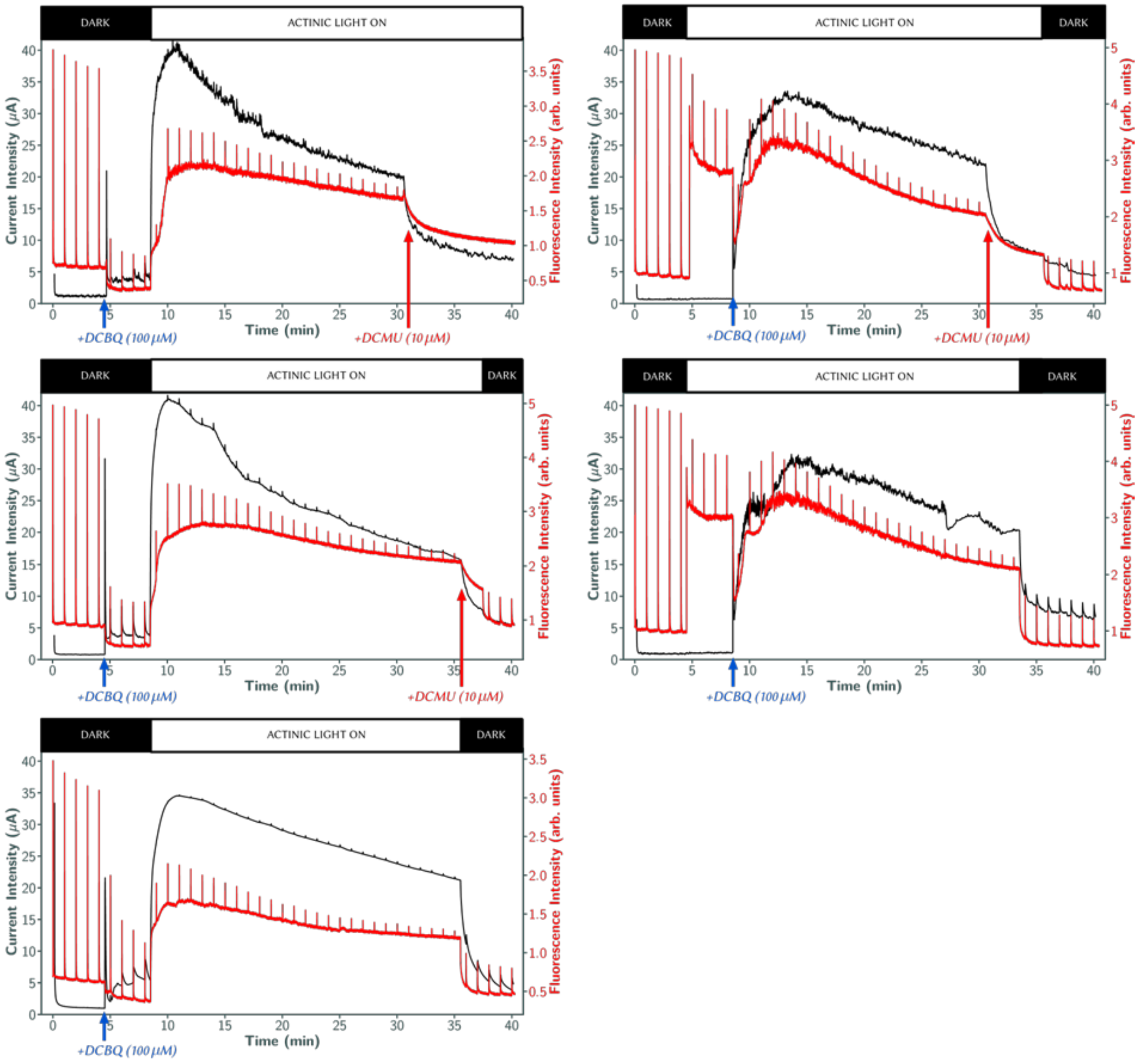


**Figure S4**. Replicates of the experiments shown in Figure 4 of the main article. Left column: 3 replicates of the experiment shown Fig.4A. Right column: 2 replicates of the experiment shown Fig.4B. In total 7 similar experiments were performed with a maximum current intensity varying between 22 µA and 42 µA (mean ± sem = 33 ± 3).
